# Topological changes in telechelic micelles: flowers versus stars

**DOI:** 10.1101/2021.08.13.455870

**Authors:** Vladimir A. Baulin

## Abstract

Micellization and morphology of spherical telechelic micelles formed by tri-block copolymers with short solvophobic end blocks at low concentrations is discussed within scaling argumentsand Single Chain Mean Field Theory (SCMFT). In ultra-dilute regime, individual telechelic polymer chains can exist in solution in two distinct states: open linear chain conformation with two free ends and closed loop conformation, when two ends are connected by the effective attraction between two solvophobic ends. At concentrations below gelation point, closed loops tend to form micelles comprised mostly of loops in flower-like micelles, while linear polymers in open conformations tend to form star-shaped aggregates with one hydrophobic dangling end. Resulting two kinds of micelles have remarkably different topology and dimensions, while the transition between them can be driven by the entropy, namely conformation changes between domination of the looped and linear conformations. The transition between two types of micelles lies in a narrow interaction parameters range. Thus, these topological micelles are very sensitive to the changes in external environment and they can serve as a very sensitive stimuli-responsive smart materials.

**Graphical TOC Entry:** 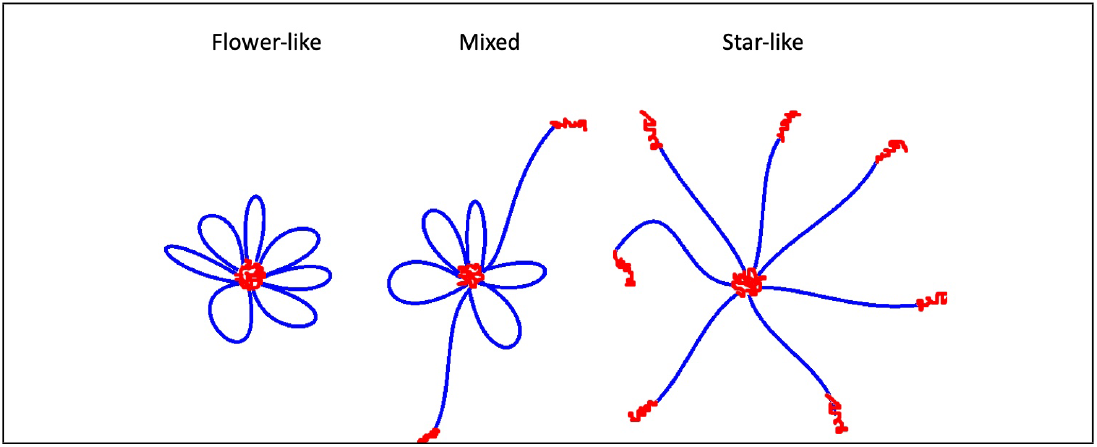

## Introduction

Stimuli-responsive smart polymers are very diverse and count a wide range of temperature and pH responsive block copolymers, hydrogels, surface-active, salt and external field responsive polymers.^1^ The sensitivity to external environments is achieved by rapid and significant conformational changes, manifested either by swelling-contraction, or internal state switches.^2^ Among stimuli-responsive polymers, a special group of telechelic associative polymers exhibit a particularly rich self-assembly behavior: with a long hydrophilic central part and short associating blocks at both ends, telechelics form diverse self-assembled aggregates^3^ ranging from single chain nanoparticles, various types of micelles and clusters, and fully connected hydrogels comprised of micelles connected with bridges.^4–6^ Such flexibility and adaptability of telechelics allow them to modulate their properties in response to external stimuli forming building blocks of architecture-transformable polymers,^7^ where the transition between different forms of architecture is purely topological, *i.e*. can be realized without breaking chemical bonds. Since these transitions between topological states are entropic in nature, the energy cost for such changes is minimal and thus, these materials can be highly sensitive to environmental changes which allows to use them as environmental sensors: dilute solution of individual flowers may abruptly change to star-shaped micelles that start to aggregate in clustered micelles. ^6^

Since the resulting telechelic micelles are of a nanometer size, only few experimental techniques may give enough spatial resolution to assess their topological state: small-angle neutron scattering (SANS) and small-angle X-ray (SAXS). A structural study of telechelic poly(ethylene oxide) (PEO) chains in a dilute regime^6^ has confirmed the formation of flower-like micelles for sufficiently high fraction of alkyl length of hydrophobic end blocks, leading to strong attraction of the hydrophobic groups and formation of clustered micelles with the presence of free dangling ends, when this attraction is reduced by reducing the alkyl length. In turn, a study^8^ of the molecular exchange kinetics between micelles suggests a multi-step process of liberating the ends. However, topological transition between individual flower-like and star-like micelles in an extremely dilute regime is expected,^6^ but has not been yet studied systematically. Individual star-shaped micelles, although not stable at contact between each other, should be observed at extremely low concentrations. In this respect, this question is similar to observation of thermodynamically stable individual polymer globule in bad solvent, which was observed at sufficiently low concentrations.^9^ In turn, environmentally induced switching between types of micelles, from flower shape to star shape, may trigger intra-micelle aggregation of star-shaped telechelic micelles and gel formation,^4,10^ leading to macroscopic changes in the solution. Thus, such conformational transitions in micelles in a very dilute regime due to changes of topology may lead to new switchable devices and sensors.

## Telechelic micelle morphologies: scaling arguments

In this section we consider geometrical arguments and scaling relations describing three morphological types of individual telechelic micelles and transitions between them due to concentration change.

The simplest structure of a telechelic polymer is a tri-block copolymer of type B-A-B, where a long hydrophilic block A is in the middle, and two short hydrophobic blocks are at the ends of a linear chain (see Figure 1). A single telechelic polymer in a solution at extra dilute concentration can be in two conformational states, a free linear chain with separated end blocks (open conformation) and a loop with aggregated end blocks (closed conformation). The conformational state of single polymers is determined by the hydrophobicity of the end blocks or specific reversible interactions^11^ or even by topological restrictions in rotaxanes.^12^

**Figure 1:**
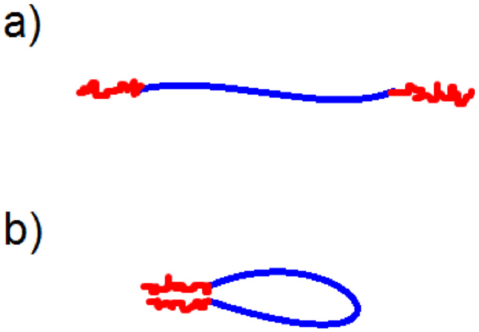
Two distinct conformations of telechelic unimers: a) hydrophobic blocks (red) are separated and b) hydrophobic blocks are reversibly connected and the hydrophilic block (blue) forms a loop.

We assume spherical aggregates of finite size, consisting of a core, formed by hydrophobic blocks and a corona, formed by hydrophilic blocks in a very dilute regime, where the micelles form finite-size aggregates and the concentration is not enough to form intra-molecular aggregates and gels. If all end blocks are in the core of the micelle, the hydrophilic blocks form loops around the core in a flower-like camomile micelle, Figure 2. If one end disconnects from the core, one of the loops transforms into a free end and the micelle has a mixed composition consisting of loops and free ends. If all loops are converted into free ends, the micelle is a star-like and resemble an aster.

**Figure 2:**
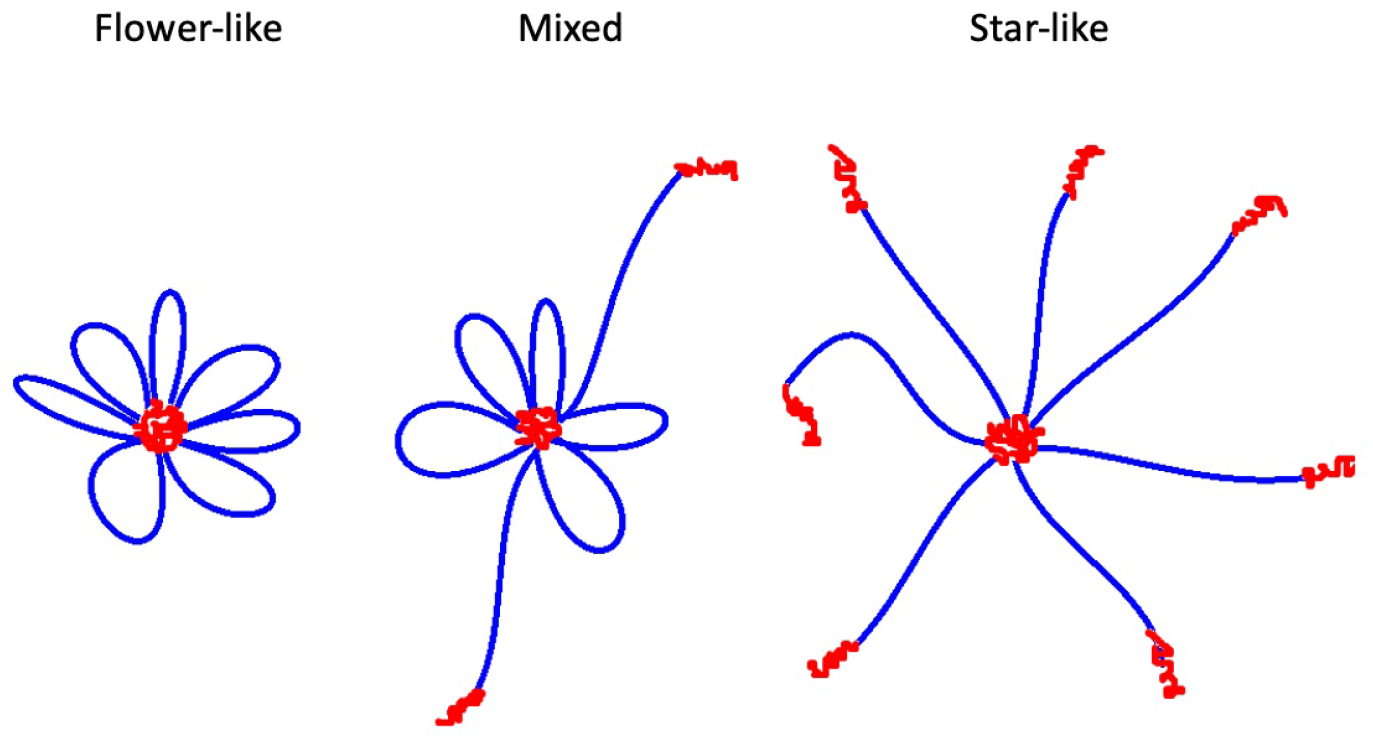
Possible structures of individual tri-block copolymer micelles: all loops are closed (flower-like or ‘‘camomile” micelle), mixture of loops and free ends (mixed micelle) and all loops are opened (a star-like or “aster” micelle).

To count conformations of such a micelle with arbitrary number of loops and free ends, we use scaling arguments originating from the renormalization group theory for polymers of arbitrary topology.^13^ In particular, we use the results obtained by Duplantier^14,15^ for the partition function of a polymer network of arbitrary topology. The partition function of the corona of the micelle has a superscaling form 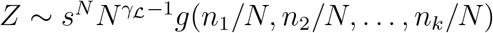, where *s* is a non-universal geometrical constant corresponding to chemical nature of the polymers, N is the total length of a network, *n*_1_, *n*_2_,…,*n_k_* are the lengths between the vertices and the ends, such that ∑*n_j_* = *N*, and *g* is a scaling function, which can be found by crosslinking different scaling regimes, when some of its arguments go either to 0 or to 1. In this expression 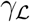 is the universal exponent which is determined only by the topology of the system and given by^15^

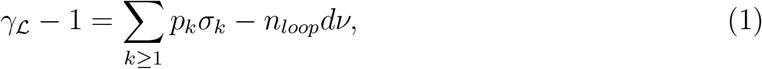

where *n_loop_* is the number of independent loops, *p_k_* is the number of vertices with *k*-legs, *d* is the dimension of space, *ν* is the Flory exponent, associated with the radius of gyration of a self-avoiding linear chain, *σ_k_* is the exponent corresponding to a vertice with *k*-legs. The universal exponents *ν* and *σ_k_* are known exactly for *d* = 2 and *d* ≥ 4 and numerically for *d* = 3. In the following we use the values of σ_k_ obtained from the simulation results for critical exponents of star polymers.^16^

According to the general expression (1), the partition function of the corona of a micelle formed by *p* tri-block copolymers, can be written in the form

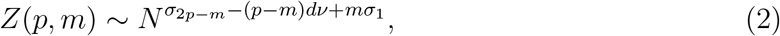

where *m* ⩽ *p* is the number of copolymers connected to the core only by one end while the rest *p* – *m* copolymers are connected to the core by both ends and form *p* – *m* loops. This expression in spirit is similar to micelles of sliding polymers, where the number of free ends and the length of the polymers in the corona are annealed.^17^ The free energy of the corona in terms of *k_B_T*, where *k_B_* is the Boltzmann constant, is given by

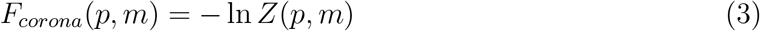

The corresponding core contribution *F_core_*(*p*, *m*) has two terms, the attraction energy of the central core and the cores of disconnected blocks

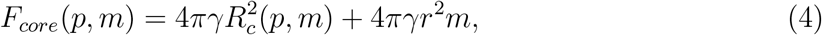

where *γ* is the surface tension between the core and the solvent, 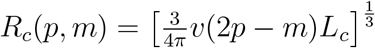 is the radius of the central core and 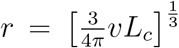 is the radius of the cores formed by disconnected blocks. Here *v* is the volume of a monomer, *L_c_* is the length of the hydrophobic block. The blocks can move inside the core and the corresponding entropy is given by

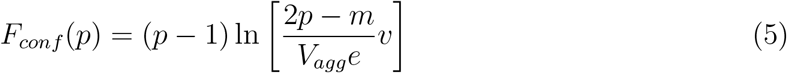

where 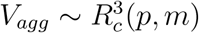. Combining all terms together, the free energy of a micelle composed of *p* tri-blocks and containing *m* free ends yields in the form

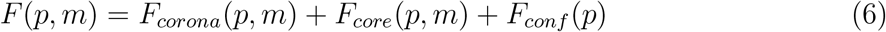

Note, that this expression is also valid for free tri-block copolymers, *p* = 1. Two configurational states depicted in Figure 1 correspond to *m* = 1 and *m* = 0. Thus, we can use *F*(1,0) or *F*(1,1) as a reference state when calculating the free energy of micellization.

The free energy of the solution containing all types of micelles is

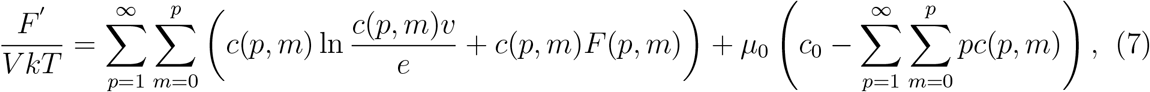

where *c*(*m*, *p*) is the number concentration of micelles composed of *p* tri-block copolymers and containing *m* free ends, *V* is the volume of the system. The last term fixes the total concentration of copolymers in the system c_0_, and the corresponding Lagrange multiplier *μ*_0_. Minimization of this free energy with respect to *c*(*m*, *p*) gives the equilibrium distribution of the micelles with respect to looped conformation of a tri-block

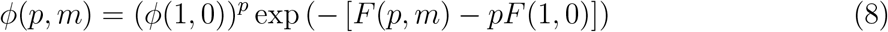

or a similar expression with a linear chain conformation as a reference state,

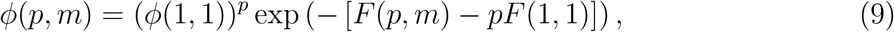

where we denote *ϕ*(*p*, *m*) = *vc*(*p*, *m*). This also means that the two concentrations *ϕ*(1,0) and *ϕ*(1,1) are related via the following expression

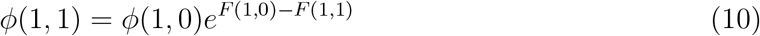

The equilibrium distribution function of micelles is complemented by the mass conservation law

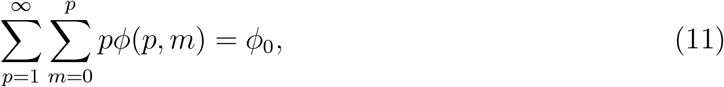

with *ϕ*_0_ = *vc*_0_. It is convenient to study the micellization process with the help of the following grand canonical function,

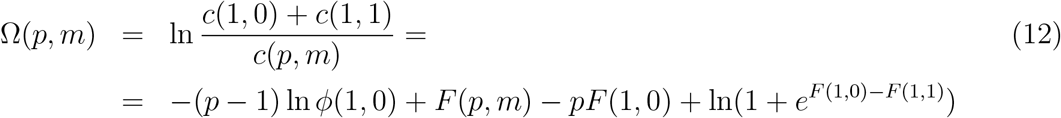

The mass action model for the equilibrium between aggregates defines the condition for the cmc, which is obtained by 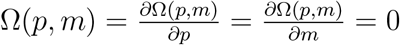, and in general it represents a line in (p,m) space. Graphically it corresponds to the minimum of the curve touching *x*-axis. If the individual chains in the solution are in closed loops conformations, the cmc associated with closed loop conformation appears first and then second, the cmc corresponding to star-like micelles. If the chains in solution are in open conformations, the flower-like micellization minimum lies above the star-like micellization and the system has only cmc, corresponding to stars. The minimum of Ω coincides with the maximum of the size distribution function *c*(*p*, *m*), indicating the average size of the micelle. Ω(*p*, *m*) = 0 corresponds to an approximate balance between free chains dissolved in the solution and the chains located in the micelles.

Tri-block copolymers self-assemble into micelles of different shapes. According to eq. (10), the ratio exp(*F*(1, 0) – *F*(1, 1)) defines the fractions of the conformations of free chains in the solution. One might expect, that this ratio influence the shape of the micelles. Indeed, if *F*(1, 0) ≫ *F*(1, 1), most of free copolymers are in the loop conformations (1, 0), thus the formation of flower-like micelles would be more favorable than the formation of star-like micelles and *vice versa*. Since the ratio between the energies of the two conformations can be tuned, *e.g*. by the temperature, one may expect temperature induced transitions between two shapes of the micelles in a narrow range. We also note that the equilibrium aggregation number of flower-like micelles should be lower than the aggregation number of star-like micelles, since the loops create more excluded volume in the corona than the free ends. Thus, the equilibrium transitions between flower-like and star-like shapes are accompanied by the changes of the sizes of the micelles and their structure. On the other hand, higher concentrations of the copolymers favor larger sizes of micelles with more crowded corona. This results in larger sizes of micelles that can be achieved more easily with star-like micelles rather than flower-like micelles. Thus, increasing concentration induce topological transition of the polymers in the micelles from loops to open chains, undergoing conformational change of entropic nature.

When *F*(1, 0) ~ *F*(1, 1) the two shapes of micelles, flower-like and star-like, can coexist (see Figure 3). The function Ω(*p*) (13) is shown for a given concentration of copolymers. The minimum of the curve related to flower-like micelles, *m* = 0, corresponds to the equilibrium size of the micelles around *p* = 12, while the minimum of the star-like micelles is around *p* = 26, *i.e*. two times larger. This situation corresponds to a critical point of the conformational transition between two types of micelles. The distribution of sizes of micelles of different shapes, *ϕ*(*p*, *m*) (8) shows that the intermediate shapes, with mixed loops and dangling ends, are also present, but their fraction is smaller. The cumulative curve, 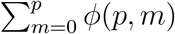, shown in grey, also shows two distinct peaks. At a low polymer concentration, the flower-like micelles form first. As the concentration increases, the star-like micelles appear and they become a dominate shape at high concentrations.

**Figure 3:**
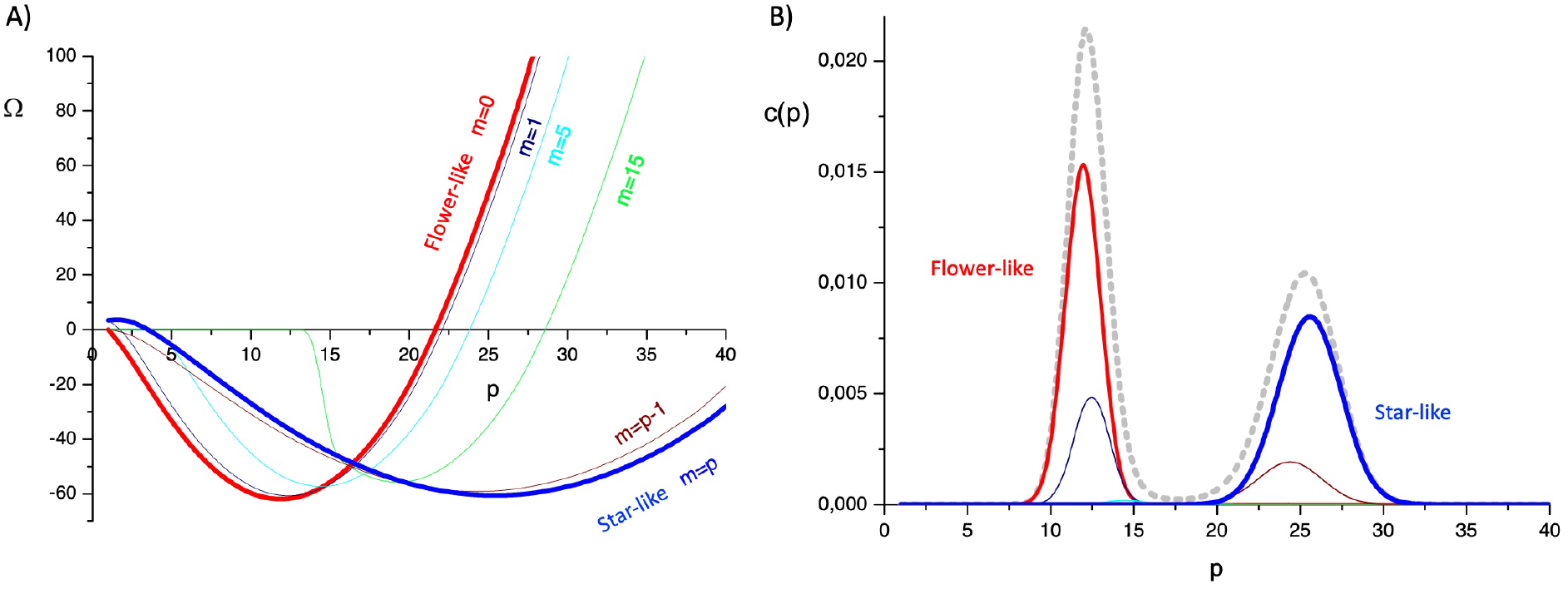
Aggregation of tri-block copolymers into micelles. A) Variation of Ω with the aggregation number *p* and B) the corresponding size distributions for *γ* = 0.57. The length of a hydrophilic block *N* = 500 and the length of the hydrophobic block *L_c_* = 50.

## Interaction-driven transitions: Single Chain Mean Field Theory

In this section, topological transitions in telechelic micelles due to environmental stimuli are considered within the Single Chain Mean Field Theory (SCMFT). The changes in the environment at a fixed concentration are modeled through the changes of the effective interaction, that controls the attraction of the telechelic ends. This could be the changes in the temperature, pH or co-solvents.

The SCMFT^18,19^ allows to get a detailed molecular structure of the micelles of self-assembled block copolymers within a mean field approximation. Here the formulation for micelles in off-lattice form is used.^20^ In this approach, the available volume is virtually divided in such a way that a micelle is placed in the center of a simulation box, while the translational entropy of the micelles of different sizes is considered through the mass action model via a corresponding translational entropy term. ^21^ A single spherical micelle placed in the center of the box has a radial symmetry and the only fields that are found self-consistently are the concentration densities of the blocks and solvent that depend only on the distance from the center of the micelle.

In general, the free energy *F* of the simulation box of volume *V*, containing p surfactants in SCMF theory is given by

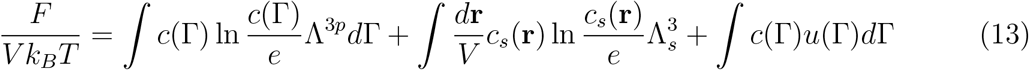

where the first term is the configurational entropy of the block copolymers in a micelle and the distribution function of conformations *c*(Γ) = *pP*(Γ)/*V* is proportional to the probability of a single molecule conformation *P*(Γ). The second term corresponds to the translational entropy of the solvent in the box with *c_s_*(**r**) being the concentration of solvent molecules, while Λ and Λ_*s*_ are the corresponding de Broglie lengths of the molecule beads and solvent molecules. Last term describes interaction of the co-polymer conformations with the fields *u*(Γ), written in a mean field form. Due to decoupling of correlations between conformations, *u*(Γ) can be written as^22,23^

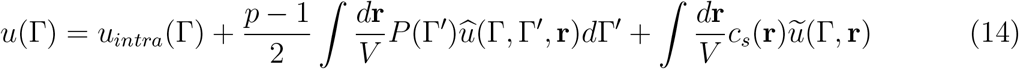

where self-interaction constant term *u_intra_*(Γ) describes interactions between blocks inside the same conformations; *û*(Γ, Γ′, **r**) describes interactions between conformations Γ and Γ′ at the position in space **r**, while *ũ*(Γ, **r**) describes the interactions with solvent.

A telechelic molecule consists of a long central block comprising of hydrophilic H beads and two very short end blocks of hydrophobic T beads, see Figure 1. Strong solvophobic interaction of the end blocks insures the properties of telechelic molecules. For the sake of simplicity, we assume only T–T attractive interactions, described by a single attractive interaction parameter *ε_TT_*, while interaction of hydrophilic beads and solvent molecules are considered only through repulsive excluded volume interactions.

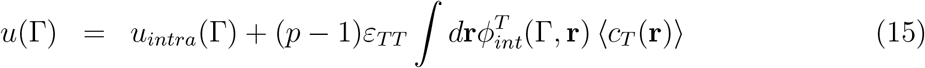

where 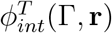 is the interaction of each conformation with the mean field of T beads in a given position in space, ⟨*c_T_*(**r**)⟩. Mean fields and the average volume fraction *ϕ*(**r**) are calculated as averages of the corresponding concentrations of each conformation, *c_T_*, *c_H_* and the excluded volume *ϕ_ex_* as

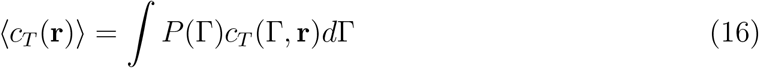

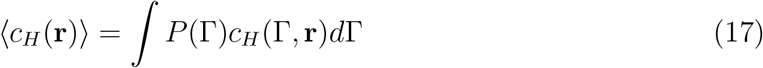

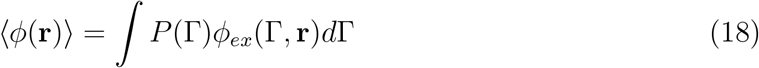

The minimization of the free energy, Eq. (13) with respect to the probabilities of the conformations *P*(Γ) subject to the incompressibility condition *ϕ_s_*(**r**) = 1 – *p* ⟨φ(**r**)⟩ gives equilibrium probabilities

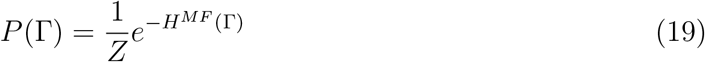

where

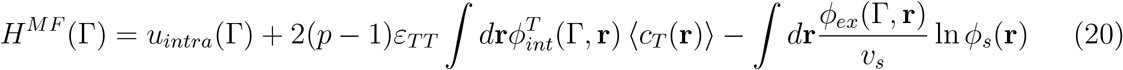

*Z* is the normalization constant, *v_s_* is the excluded volume of the solvent. Solving these nonlinear equations self-consistently, one gets the equilibrium distribution of average concentrations of the blocks as well as the most probable conformations in the equilibrium self-assembled aggregates.

Considering the size of the beads as a scale of the system, we model telechelic molecules as freely-joined chains of beads with diameter 1 and segment length 1.47. *p* telechelic molecules consisting of 20 hydrophilic beads in the central block and 1 hydrophobic bead at each end, T_1_H_20_T_1_, are placed in the square box 30 × 30 × 30, which determines the average concentration of polymers. The sampling of 10 million self-avoiding conformations is generated using Rosenbluth-Rosenbluth algorithm. ^24^ The interaction between hydrophobic beads is modeled as a potential well of width 1.61 and depth *ε_TT_*.

At relatively strong attraction of the between hydrophobic beads, *ε_TT_* = −3.8 the micelles are formed at low aggregation numbers and the most probable conformations correspond to a closed state, when two ends are connected and hydrophilic blocks form a loop. This corresponds to flower-like micelles, green curve at Figure 4A). At weaker attractions, *ε_TT_* = −1.9, telechelics assemble into star-like micelles with most probable conformations in open state, orange curve at Figure 4A). The shift of the minimum to larger aggregation numbers is due to smaller entropic repulsion of free ends compared to the loops. The volume fraction of flower-like micelle, Figure 4B) and star-like micelle, Figure 4C) also shows a structural difference between the micelles.

**Figure 4:**
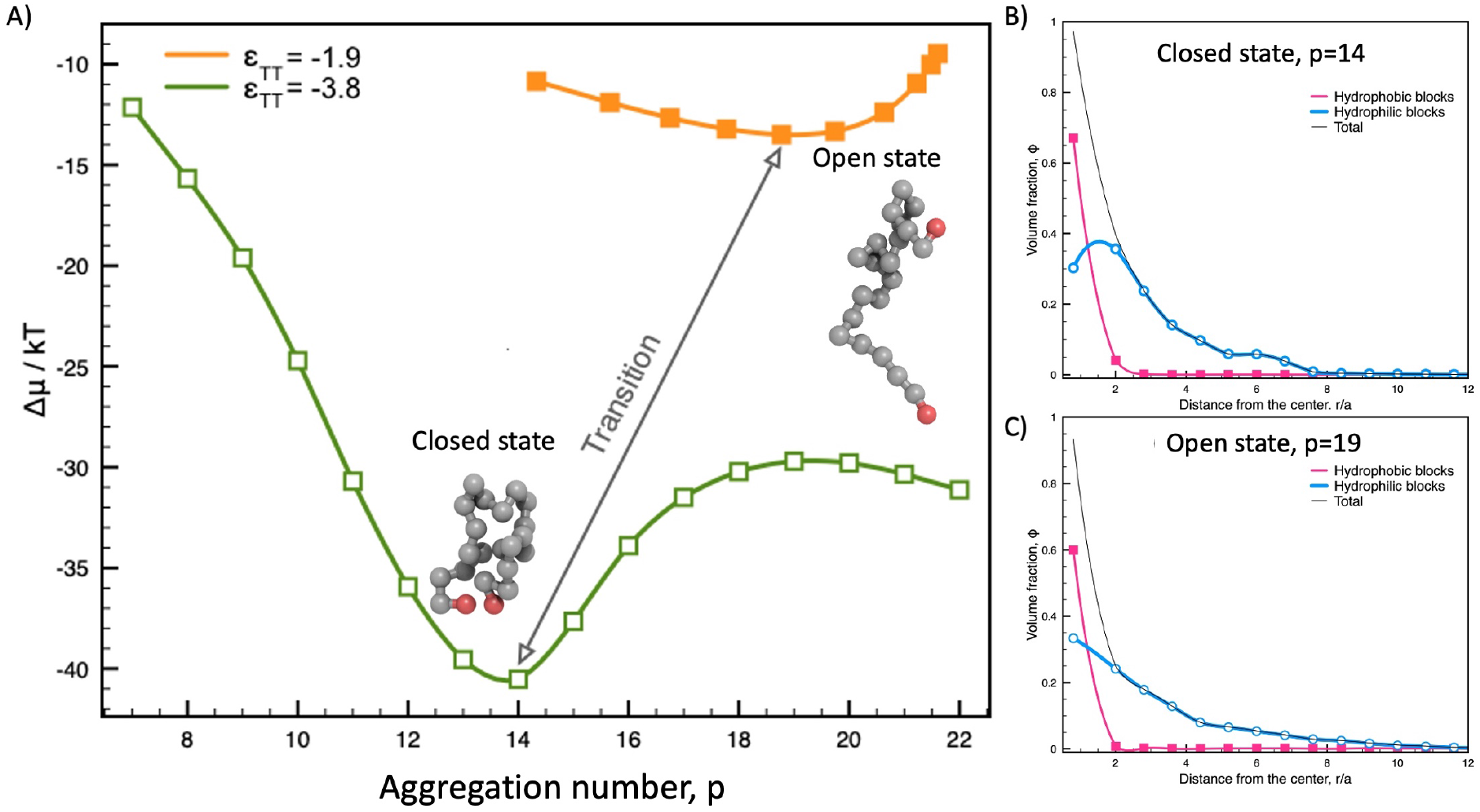
SCMF theory of telechelic micelles. A) Chemical potential difference Δ*μ* between monomers in the micelles and in the solution as a function of the aggregation number *p* for strong attraction, *ε_TT_* = −3.8, flower-like micelle, green curve, open squares, and weak attraction, *ε_TT_* = −1.9, star-like micelle, orange curve, closed squares. The corresponding to the minima volume fractions of hydrophobic blocks (red), hydrophilic blocks (blue) and the sum (grey) as a function of a distance from the center in the B) flower-like micelle, *p* =14 and C) star-like micelle, *p* = 19.

Thermodynamically stable flower-like micelles exist in a narrow parameter space: the interaction between the end groups should be relatively strong to prevent opening due to entropy of ends and the hydrophilic central block should be relatively long. Thus, small changes in the environment can provoke the transition to star-like aggregates that can further aggregate between each other, providing a tool for sensing the changes in the solvent.

## Conclusion

In conclusion, a scaling theory of conformational changes in telechelic micelles in a dilute regime demonstrates the existence of two types of micelles that can exhibit conformational transitions between them with increase of concentration. The type of a micelle (flower- or star-like) is determined by the conformation of the individual chains in the solution. If the attraction between two ends is weak, individual chains in the solution are preferentially in open conformations (free ends), which subsequently self-assemble in star-like micelles and flower-like micelles are not observed for any concentration. If the attraction between the blocks is strong enough for hydrophobic ends to form a stable closed loop conformation, individual chains in the solution are in a closed state (a loop) and subsequently self-assemble into flower-like micelles comprising of a hydrophobic core surrounded by hydrophilic loops. With increasing concentration, the aggregation number increases until the loops are no longer favorable due to the excluded volume restrictions near the center of the micelle and the micelle transforms to a star-like micelle by a jump without intermediate states, which is typical for topological transitions.

The mean field approach within the SCMF theory also confirms the existence of flower- and star-like micelles and provides microscopic details of the structure of the micelles of both types. The changes in the environment may induce the changes in the topology of the micelle, which can undergo sharp transition from flower-like to star-like micelle. The surface properties of the two types of micelles can be very different: flower-like micelles have exclusively hydrophilic blocks in the outer shell, while star-like micelles have sticky hydrophobic blocks in the corona, which can bridge or connect with other hydrophobic objects such as *e.g*. lipid membranes. Due to the entropic nature, topological transitions are highly sensitive to external environment changes and thus can be triggered by external changes in the environment in numerous applications as sensors or stimuli-responsive polymers.

## Acknowledgement

This publication is a part of the project I+D+i: PID2020-114347RB-C33, financed by MCIN/ AEI 10.13039/501100011033.

The manuscript was deposited at bioRxiv doi:10.1101/2021.08.13.455870

